# Meta Learning Improves Robustness and Performance in Machine Learning-Guided Protein Engineering

**DOI:** 10.1101/2023.01.30.526201

**Authors:** Mason Minot, Sai T. Reddy

## Abstract

Machine learning-guided protein engineering continues to rapidly progress, however, collecting large, well-labeled data sets remains time and resource intensive. Directed evolution and protein engineering studies often require extensive experimental processes to eliminate noise and fully label high-throughput protein sequence-function data. Meta learning methods established in other fields (e.g. computer vision and natural language processing) have proven effective in learning from noisy data, given the availability of a small data set with trusted labels and thus could be applied for protein engineering. Here, we generate yeast display antibody mutagenesis libraries and screen them for target antigen binding followed by deep sequencing. Meta learning approaches are able to learn under high synthetic and experimental noise as well as in under labeled data settings, typically outperforming baselines significantly and often requiring a fraction of the training data. Thus, we demonstrate meta learning may expedite and improve machine learning-guided protein engineering.

**Availability and implementation:** The code used in this study is publicly available at https://github.com/LSSI-ETH/meta-learning-for-protein-engineering.

**Graphical Abstract:** 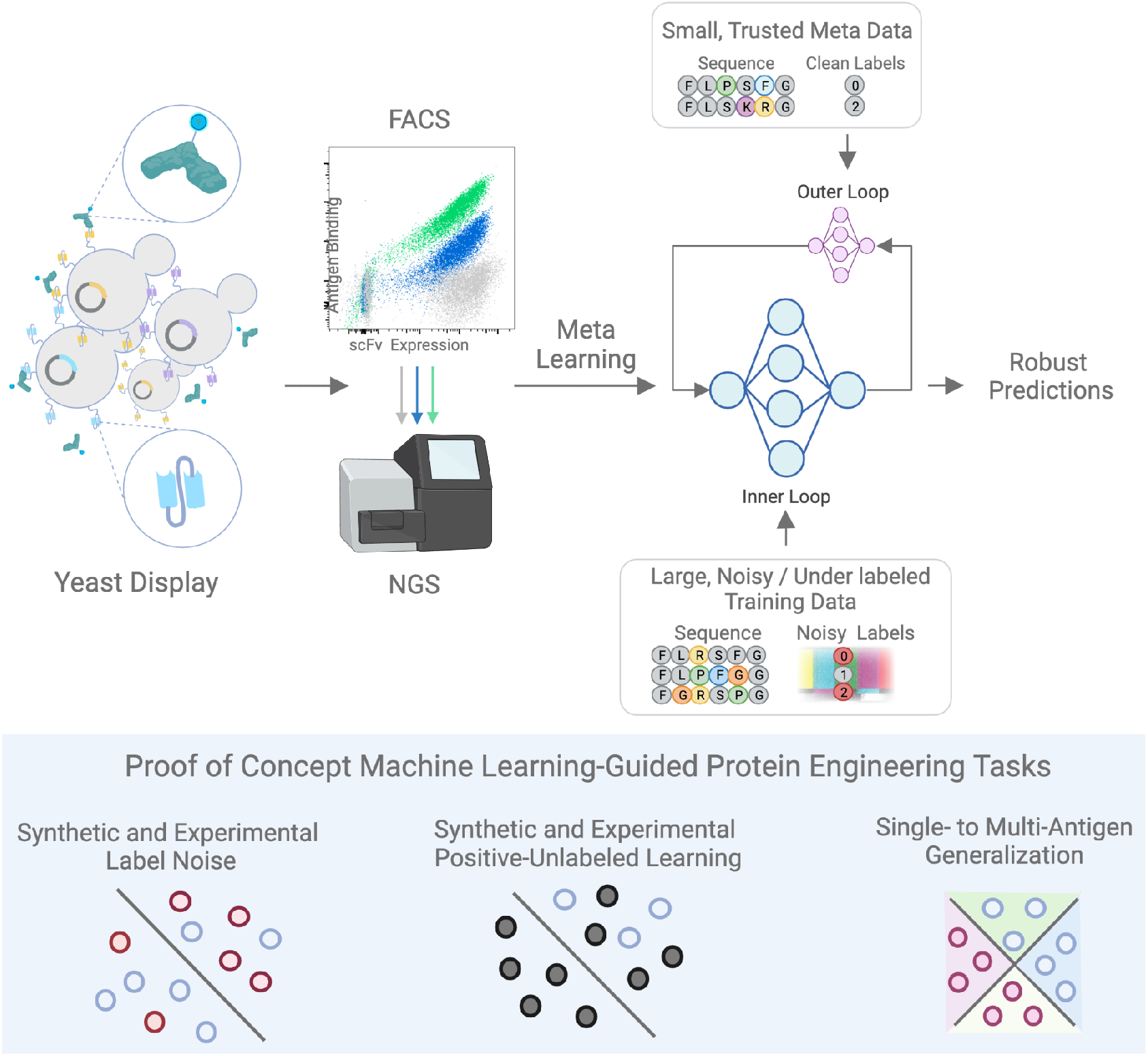

## Introduction

Machine learning-guided protein engineering is an emerging field that integrates a number of tools and techniques including deep sequencing, DNA synthesis, high-throughput screening, structural modeling and computational algorithms (Yang *et al*., 2019). A number of recent studies are demonstrating the potential of this field, for example machine learning-guided structural mutagenesis was used to improve the enantioselectivity and thermostability of a hydrolase enzyme used for degrading plastic waste (Lu *et al*., 2022). Machine learning has been used in antibody engineering as well, such as predicting therapeutic antibody specificity for developability assessment (Mason *et al*., 2021), off-target or polyspecific binding (Makowski *et al*., 2022; Saksena *et al*., 2022), and predicting escape of viral variants (e.g., SARS-CoV-2) to antibody drug candidates (Taft *et al*., 2022). Furthermore, the de novo design of synthetic proteins via generative modeling has leveraged data driven deep learning to outperform physically based design methods on a variety of tasks including improving protein expression, stability, and ligand binding (Dauparas *et al*., 2022; Wicky *et al*., 2022; Shin *et al*., 2021).

The application of machine learning in directed evolution is particularly well suited as high-throughput screening systems (e.g., phage or yeast display) (Clackson *et al*., 1991; Boder and Wittrup, 1997) link protein sequence variants (genotype) with functional properties (phenotype). This enables the use of supervised machine learning methods, which rely on labeled data for training models. For example, display libraries of antibody variants can be screened for binding and non-binding to target antigen by fluorescence activated cell sorting (FACS) followed by deep sequencing, resulting in labeled data sets for supervised machine learning (Makowski *et al*., 2022; Bryant *et al*., 2021; Mason *et al*., 2021). However, collecting high-quality, large, and well-labeled data sets requires extensive experimental workflows and can represent a bottleneck. A number of studies have set out to reduce experimental burden, for example through smart training set design (Wittmann *et al*., 2021), the use of large protein language models trained on massive protein databases (Ofer *et al*., 2021; Rao *et al*., 2021; Biswas *et al*., 2021), and data augmentation techniques (Minot and Reddy, 2022; Han *et al*., 2019; Li and Zhang, 2021). While these approaches serve to maximize the use of available data, they may not generalize well without high-quality, fully labeled data.

High-throughput screening experiments often require multiple rounds of enrichment (e.g., FACS) to eliminate noisy (false positives/false negatives) or under labeled data (Harper *et al*., 2001). Under labeled data can include unlabeled samples or those with missing labels, e.g possessing binding or non-binding labels to only a subset of the targets of interest in a multi-target classification setting. Previous approaches to address noise in biological assays include Naïve Bayes classifiers (Glick *et al*., 2004) and Binary Kernel Discrimination (Harper *et al*., 2001) applied to small molecule screens for pharmaceutical development. However, such methods are lacking in protein engineering and predate recent advances in deep learning. Furthermore, while high-throughput screening assays can yield protein variants enriched for a desired property, obtaining true negatives necessary for supervised machine learning may be intractable in some cases (e.g. growth-based selections). In these instances, experimental endpoints typically result in deep sequencing of the positive population and the unlabeled library prior to screening (Song *et al*., 2021). This unlabeled population contains both positives and negatives, thereby complicating supervised learning approaches. Thus, machine learning-guided protein engineering could benefit from methods applicable to noisy and under labeled data.

Here, we propose meta learning, also known as ‘learning to learn,’ as a method to improve the robustness and performance of machine learning-guided protein engineering. Although the field of meta learning is large, we focus on recent advances for noisy data (Ren *et al*., 2018; Zheng *et al*., 2021). Experimentally, these approaches train with a large set of noisy data and use a small data set of clean, trusted examples, referred to as a meta set. Computationally, the framework typically consists of nested optimization loops, with the inner loop trained on noisy data and the outer loop guiding the inner loop and preventing fitting to the noise using the meta set. As a proof-of-concept we generate multiple yeast display antibody mutagenesis libraries and screen them for target antigen binding followed by deep sequencing. Several machine learning models and classification tasks are evaluated in the context of noisy or under labeled data. These tasks include synthetic and experimental label noise, learning from synthetic and experimental positive and unlabeled (PU) data, and multi-antigen binding classification from largely single-antigen data. Meta learning regularly results in impressive performance improvements over baseline models even with very high label noise, and in low data regimes. Taken together, these results serve to demonstrate the power and potential of meta learning in machine learning-guided protein engineering.

## Results

### Design and experimental screening of antibody mutagenesis libraries

To establish a meta learning approach for protein engineering, we first designed yeast display antibody libraries (single-chain fragment variable, scFv) (Boder and Wittrup, 1997). The parental antibody clones 4D5 and 5A12 were chosen as they have extensive background information, including antibody-antigen co-crystal structures (Koenig *et al*., 2015; Cho *et al*., 2003) and antigen-binding deep mutational scanning (DMS) data (Koenig *et al*., 2015; Fowler and Fields, 2014; Mason *et al*., 2021). 4D5 is a scFv derived from the therapeutic antibody Trastuzumab and binds to the human epidermal growth factor receptor 2 (HER2) (Carter *et al*., 1992). 5A12 has the unique property of multi-specificity, as it has high affinity towards two structurally unrelated antigens, human vascular endothelial growth factor (VEGF) and human angiopoietin-2 (Ang2) (Koenig *et al*., 2015). To maximize the number of function-retaining mutations, two complementarity determining region (CDR) loops per scFv were chosen for modification based on literature data: the CDRH3 and CDRL3 in 4D5 and the CDRH2 and CDRL1 in 5A12. A total of nine amino acids were targeted for mutagenesis in each antibody, where crystal structures demonstrated interactions of these positions with the target antigens (Fig. 1 B,E,H). The yeast display mutagenesis libraries, along with parental clones as controls were screened with cognate fluorescently-labeled antigens and sorted by FACS to obtain multiple populations (Fig. 1 C,F,I and Fig. S1, S2, S3). The 4D5 library was sorted into three separate populations based on HER2 antigen binding strength (High-, Low-, and Non-binding fluorescence intensity gates) (Fig. 1 C, Fig. S1). The 5A12 library was screened in a binary manner for High- or Low/Non-binding to VEGF or Ang2 (Fig. 1 F,I and Fig. S2, S3). Following sorting, targeted deep sequencing was performed on the scFv variants for each population and diversity is visualized through protein sequence logo plots (Fig. 1 D,G,J).

**Fig. 1.**
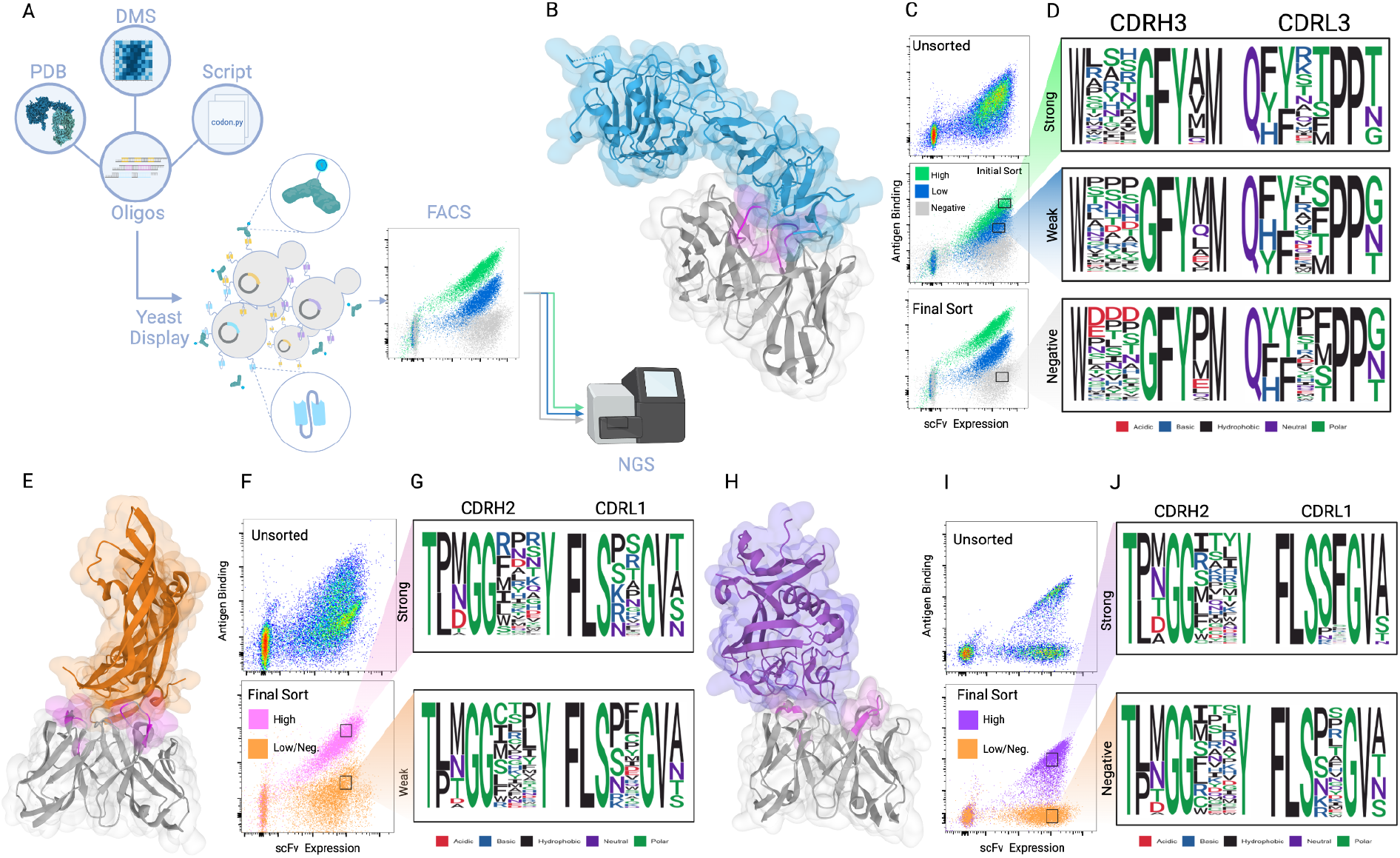
Design and screening of yeast display antibody (scFv) libraries. **(A)** Workflow overview: Degenerate oligonucleotides are computationally designed based on previously published deep mutational scanning data (DMS) (Mason *et al*., 2021; Koenig *et al*., 2015) and crystal structures (PDB: 1N8Z, 4ZFF, 4ZFG), transformed into yeast, and screened by FACS for binding to antigen. The sorted populations are then used for targeted deep sequencing of the antibody variable regions. **(B)** The crystal structure of 4D5 scFv (grey) in complex with HER2 (blue) (PDB: 1N8Z). The amino acid positions of 4D5 targeted for combinatorial mutagenesis are highlighted in pink. **(C)** The 4D5 scFv library is screened for binding to HER2 antigen by FACS; dot plots show the library is fractionated into HER2-binding populations (High-, Low-, or Non-binding) based on fluorescence intensity. Multiple rounds of FACS are performed (Fig. S1). **(D)** Protein sequence logo plots represent the mutagenesis regions of the 4D5 scFV and are derived from deep sequencing of the various HER2-binding populations. **(E)** The crystal structure of 5A12 scFv (grey) in complex with VEGF antigen (orange) (PDB: 4ZFF). The amino acid positions of 5A12 targeted for combinatorial mutagenesis are highlighted in pink **(F)** The 5A12 scFv library is screened for binding to VEGF antigen by FACS; dot plots show the library is fractionated into VEGF-binding populations (High- or Low/Non-binding) based on fluorescence intensity. Multiple rounds of FACS are performed (Fig. S2). **(G)** Protein sequence logo plots represent the mutagenesis regions of the 5A12 scFV and are derived from deep sequencing of the various VEGF-binding populations. **(H)** The crystal structure of 5A12 scFv (grey) in complex with Ang2 antigen (orange) (PDB: 4ZFG). The amino acid positions of 5A12 targeted for combinatorial mutagenesis are highlighted in pink **(I)** The 5A12 scFv library is screened for binding to VEGF antigen by FACS; dot plots show the library is fractionated into Ang2-binding populations (High- or Low/Non-binding) based on fluorescence intensity. Multiple rounds of FACS are performed (Fig. S3). **(J)** Protein sequence logo plots represent the mutagenesis regions of the 5A12 scFV and are derived from deep sequencing of the various Ang2-binding populations.

### Establishing meta learning workflows

The results of antibody library screening experiments and deep sequencing data were used to determine if meta learning could augment machine learning models in the prediction of sequence-function relationships based on a series of proof-of-concept tasks. The tasks were constructed to assess learning from data with synthetic or experimental noise, positive (antigen-binding) and unlabeled (containing both binding and non-binding) classification, and multi-antigen classification. Each data set was divided into training, meta, and test sets. For each task, training and testing sets were separated via edit distance from the consensus sequence of the full data set (Methods). This approach is common in machine learning-guided protein engineering as it resembles real-world workflows, in which models are trained with a limited number of mutations and used to extrapolate to a larger sequence space (Dallago *et al*., 2021). In each workflow, the large training sets consisted of antibody sequences and labels that are noisy (false positives and false negatives) or under labeled (partially labeled, e.g. for one out of two antigens, or completely unlabeled), while the meta sets are small and clean (experimentally validated true positive and true negative labels) antibody sequences under the edit distance cutoff. The test sets are large, clean, and fully labeled sequences above the edit distance cutoff (Table S5).

#### Machine Learning Models and Baselines

All training, meta, and test data of antibody protein sequences were categorically or one-hot encoded (Methods) and used in base models with different training algorithms. The main base models used for this study include a convolutional neural network (CNN) (LeCun and Bengio, 1998) and a transformer (Vaswani *et al*., 2017), which have demonstrated reliable performance in protein engineering studies (Mason *et al*., 2021; Rives *et al*., 2021; Wu *et al*., 2020). As the purpose of this work is not to develop the best model for each task, but to investigate the utility of meta learning, simple sequence encoding methods were used, minimal hyperparameter optimization was performed, and the same hyperparameters are used across all tasks (see Methods). Two baseline training algorithms were used in accordance with common practice: ‘Fine-Tune’ (FT) and ‘MetaSet-Only’ (MSO) (Ren *et al*., 2018; Shu *et al*., 2019). The Fine-Tune baseline applies a traditional training protocol to the noisy training data, however, following the completion of training, the model is then fine-tuned on the clean meta set. Fine-Tune is considered an appropriate baseline in noisy meta learning as it allows both meta and non-meta algorithms access to the same clean and noisy data. The MetaSet-Only baseline uses a traditional training protocol to train on the small, clean meta set only, without training on noisy data and typically results in poor performance due to extensive overfitting to the small data (Table S6). As additional benchmark controls for each task, we tested traditional machine learning models (Random Forest, Linear Support Vector Classifier, and Naive Bayes) (Fig. S4). These control models consistently underperformed meta learning approaches and most other baselines. Finally, we tested the meta learning algorithms on Logistic Regression and a standard Multilayer Perceptron, which yielded noteworthy results in some cases, but were typically outperformed by the meta learning-supported CNN and transformer (Fig. S5). Standard Random Forest, Linear Support Vector Classifier, and Naive Bayes classifiers are not compatible with the meta learning algorithms as they are not trained using stochastic gradient descent.

#### Meta Learning Algorithms

We chose two orthogonal, representative meta learning approaches to learn from noisy data: learning to reweight (L2RW) and meta label correction (MLC) (Ren *et al*., 2018; Zheng *et al*., 2021). Both approaches are nested optimization algorithms consisting of an outer loop meta learner which guides the training of a traditional classifier in the inner loop (Fig. 2, adapted from (Ren *et al*., 2018; Zheng *et al*., 2021)). Both inner and outer loops are trained via stochastic gradient descent and the resulting outer loop training process can be thought of as gradient descent on the gradient descent of the inner loop (‘learning to learn’). However, the two algorithms differ in their meta learning objective.

**Fig. 2.**
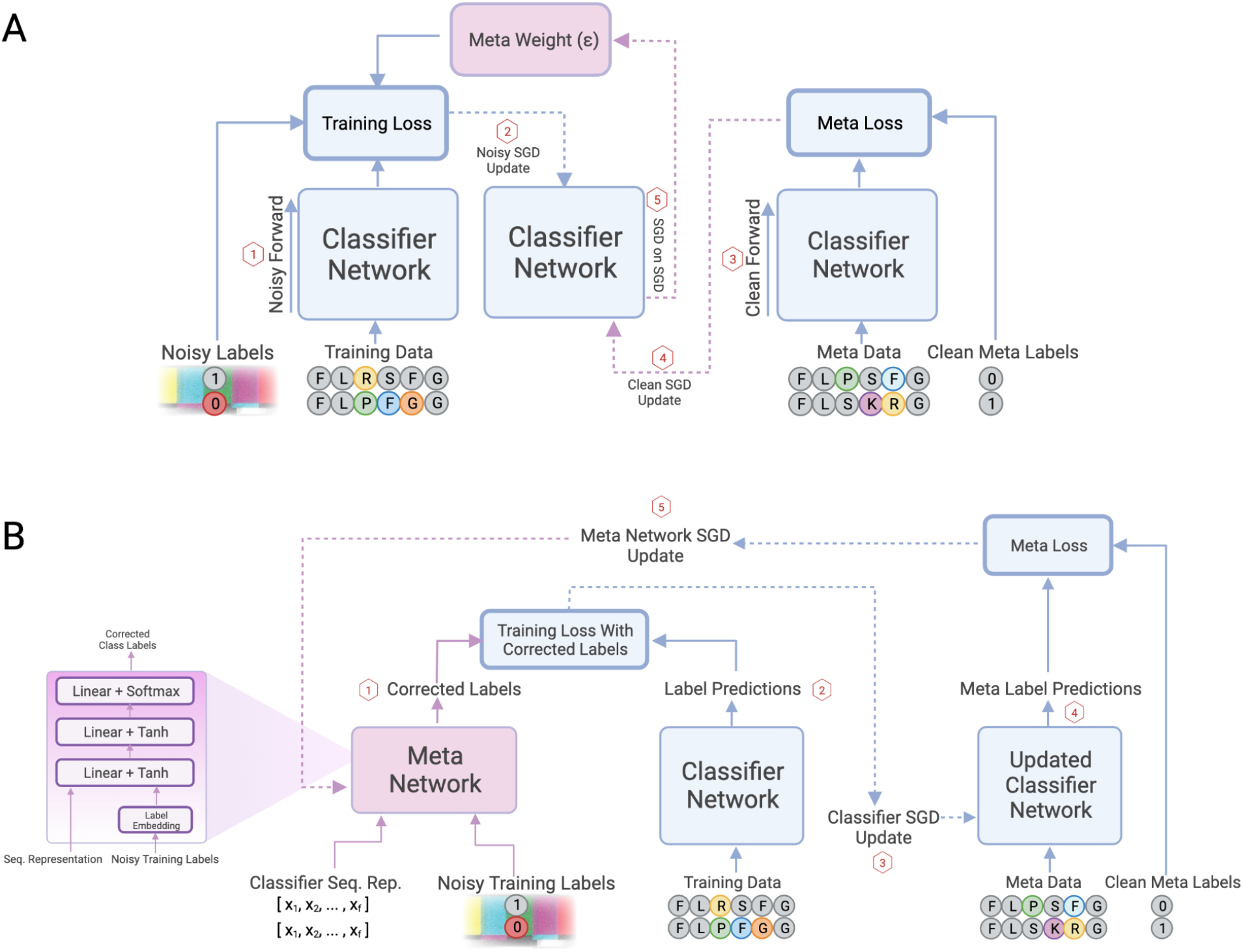
Meta learning algorithms for prediction of antigen binding trained with deep sequencing of yeast display antibody (scFv) libraries and noisy binding classification labels. **A)** Schematic representation of L2RW (Ren *et al*., 2018). The inner loop is trained on protein sequences with noisy labels. The outer loop adaptively assigns importance weights to training examples with the goal of down-weighting noisy examples, thereby scaling the loss and therefore the magnitude of the weight updates during gradient descent. The sequence of operations for training with a single mini-batch is numbered in red. Solid lines indicate forward passes. Dashed lines indicate backpropagation. Blue lines correspond to the classifier network (CNN or transformer). Purple lines correspond to the meta learner. **B)** Schematic representation of MLC (Zheng *et al*., 2021). A meta label correction network to identify and correct noisy labels is linked with a standard classifier (CNN or transformer) trained on protein sequences. The inner loop gradient descent updates the standard classifier weights from noisy data with corrected labels. The outer loop gradient descent uses the clean meta set to guide training. The sequence of operations for training with a single mini-batch is numbered in red. Solid lines indicate forward passes. Dashed lines indicate backpropagation. Blue lines correspond to the classifier network (CNN or transformer). Purple lines correspond to the meta learner.

LR2W is based on the observation that noisy examples are often difficult for the network to classify and therefore produce a greater loss (Ren *et al*., 2018). Using a small, clean meta set as guidance, the outer loop of L2RW adaptively assigns importance weights to training examples. The importance weights scale the loss, thereby scaling the magnitude of the weight updates during gradient descent. In reducing the gradient update from suspected noisy training samples, the algorithm attempts to prevent the network from fitting to noise.

The second algorithm, MLC, links a standard classifier with a meta network that attempts to identify and correct noisy labels (Zheng *et al*., 2021). The inner loop gradient descent updates the standard classifier weights from noisy data with corrected labels. The outer loop gradient descent update is performed using the clean meta set. One critique of reweighting approaches is that they may down-weight outliers or difficult training examples that may produce high loss, but are in fact, correctly labeled. In contrast, MLC seeks to avoid discarding outliers and difficult examples via its meta label correction network. This approach can in general lead to overall improved performance, but often requires larger training and meta sets.

It is worth noting that despite impressive performance in difficult learning scenarios, meta learning comes with its own challenges. For example, nested optimization requires up to three gradient descent steps per training batch, as opposed to one in standard stochastic gradient descent, thereby increasing computational cost and training time up to threefold (Fig. 2). Meta learning is also sensitive to learning rate choice and can fail to learn outside of an appropriate regime, therefore careful selection is required. However, theoretical recommendations for learning rate selection have been investigated (Shu *et al*., 2019; Ren *et al*., 2018).

### Benchmarking meta learning with synthetic noise

As a proof-of-concept, we first tested the meta learning approaches on synthetically corrupted data. Using antibody deep sequencing data from the 4D5 libraries from the final round of FACS (High-, Low-, and Non-binding to HER2), each population was assigned a class label based on its binding ability (Methods). Antibody sequences derived from this final round of FACS enrichment result in highly accurate classification labels (High-, Low-, and Non-binding to HER2) with minimal experimental noise as populations were well separated (Fig. S1). Sequence labels in the training set were then corrupted by probabilistically flipping them to another class label based on a predefined noise fraction (η) ranging from 0 to 0.9. Sequence labels in the test and meta data sets remained unaltered.

We also sought to identify the impact of training data size on model performance. This is especially relevant in an experimental context as generating data is time and resource intensive. We truncated our training set to sizes ranging from 1% to 100 % of available training data. We chose a clean meta set size of 96 antibody sequences (Methods), as it is reasonable to validate antibody binding for this many variants using experimental assays (e.g., enzyme-linked immunosorbent assays). The meta set was obtained from the final round of enrichment (FACS) and thus expected to have accurate classification labels resulting from extensive screening and good separation. Classification performance is assessed using Matthew’s Correlation Coefficient (MCC), which ranges between −1 and 1. Model performance is plotted as a function of training data size and noise fraction for the transformer and CNN base models using the baseline and meta training algorithms (Fig. 3). Baseline performance rapidly drops as noise fraction is increased, however, both L2RW and MLC retain high performance under high label noise. L2RW maintains a high performance even at low training set sizes when noise is low, but yields a lower overall performance as noise fraction, η, increases past 50 %. In contrast, MLC exhibits low performance for small training sets, but maintains a substantial 0.8 MCC overall once the training set size is increased to 20 % or more of the available data. In general, meta learning yields impressive results, even under very high label noise with small amounts of training data and a small meta set.

**Fig. 3.**
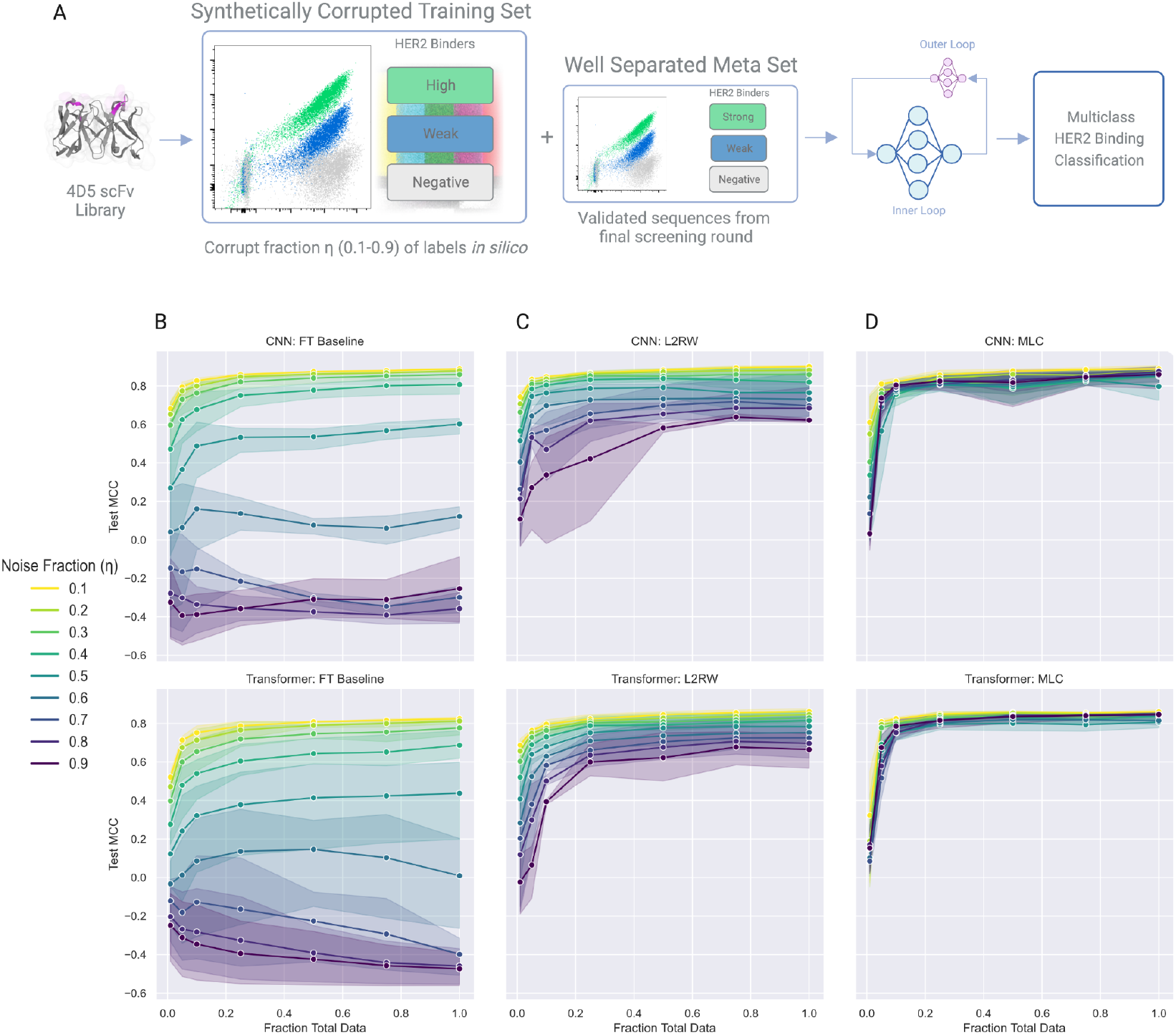
Meta learning applied to supervised machine learning models trained from yeast antibody (scFv) sequence data with synthetic label noise. **(A)** Schematic representation of machine learning task. Train, test, and meta data sets consist of deep sequencing of 4D5 libraries following the final enrichment round of FACS for binding to HER2 antigen (Fig. S1). Class labels refer to 4D5 variant populations with High-, Low-, and Non-binding to HER2 antigen. Training set labels are probabilistically corrupted according to specified noise fraction η, while meta and test set data labels remain unmodified. Models are evaluated based on multiclass predictions of 4D5 scFv variants binding to HER2 antigen. **(B, C, and D)** Meta learning and baseline machine learning model prediction performance is based on Matthew’s Correlation Coefficient (MCC) and plotted as a function of the number of training samples. FT refers to Fine-Tune Baseline. Points correspond to mean performance and shaded regions to 95 % confidence intervals across 3 random seeds. Performance curves are colored according to the training set noise fraction η. A meta data set of 96 sequence-function pairs was used for all models.

### Applying meta learning on data sets with experimental noise

After having benchmarked meta learning on synthetically generated noise, we next tested the approach on data sets with experimental noise. Screening of antibody or protein mutagenesis libraries by display platforms often requires multiple rounds of enrichment (FACS) to remove undesired mutants (e.g., false positives), thus requiring substantial time and resources. We sought to determine if meta learning could accelerate the screening process by learning from an unenriched antibody mutagenesis library, such as after only a single round of screening. Therefore the training set consists of antibody deep sequencing data from the populations (High-, Low-, and Non-binding to HER2) after only the first round of FACS. A single round of sorting shows there is overlap across these populations indicating that a substantial fraction of noise will be present (antibody sequences with misclassified labels) (Fig. 4A, S1). Following training, machine learning model performance was assessed by testing on sequences from the fourth round of enrichment, at which point the populations were well separated due to extensive screening and expected to have minimal experimental noise (Fig. 4B). As before, meta sets were derived from this fourth round of enrichment. The impact of training set size and different meta set sizes on machine learning model predictions of the test data was assessed. MCC values showed improved performance across baseline and meta algorithms as meta size is increased from 32 sequences to 864 sequences. The Fine-Tune baseline performs poorly when training data is minimal, however, MCC rises to approximately 0.8 for full training data and large meta sets.

**Fig. 4.**
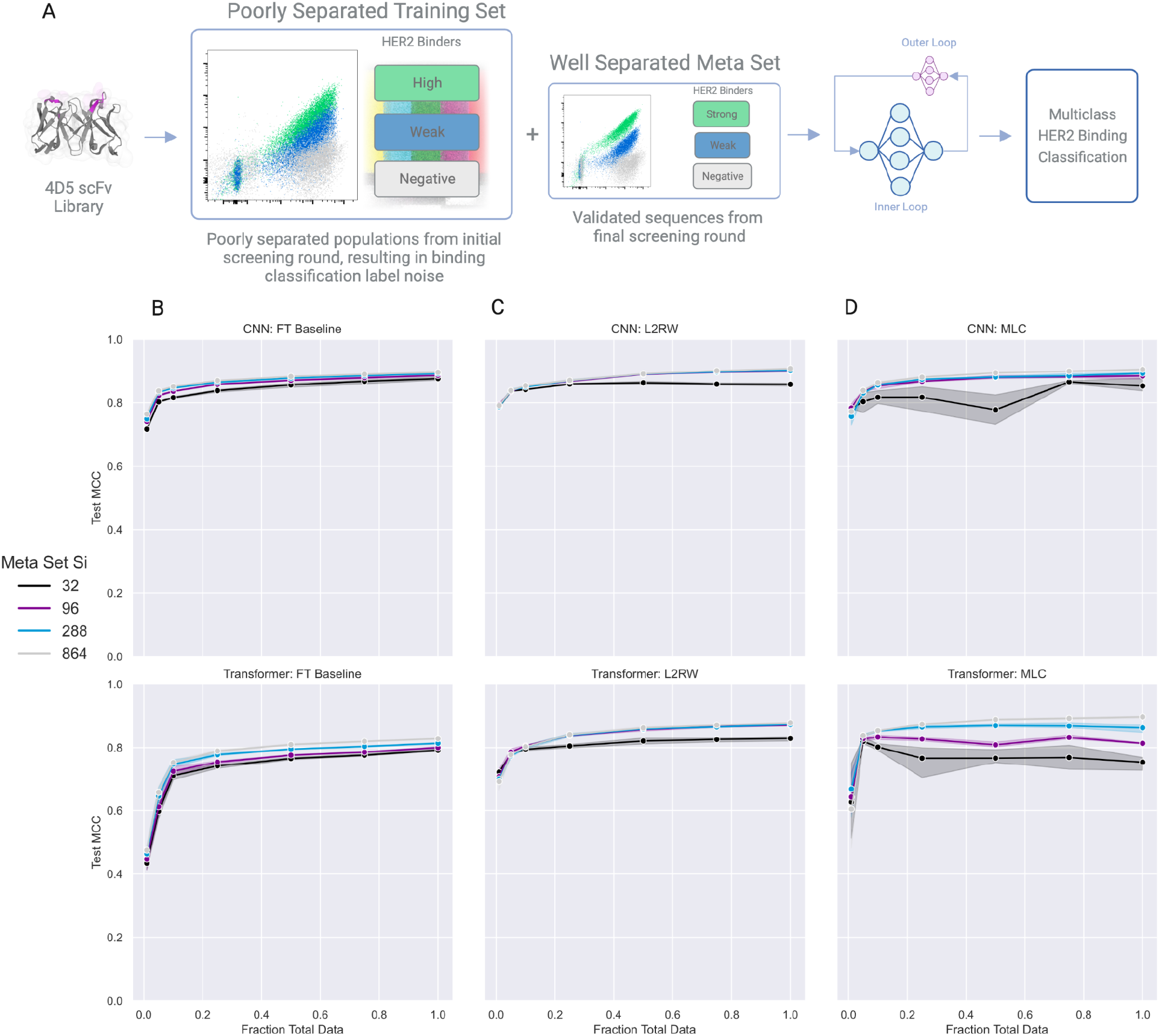
Meta learning applied to supervised machine learning models trained from yeast antibody (scFv) sequence data with experimental label noise. **(A)** Schematic representation of machine learning task. Training data sets consist of deep sequencing of 4D5 libraries from the initial round of FACS, at which point populations are poorly separated, resulting in noisy binding classification labels (Fig. S1). Test and meta data sets consist of deep sequencing of 4D5 libraries from the final round of FACS. Class labels refer to 4D5 variant populations with High-, Low-, and Non-binding binding to HER2 antigen. Models are evaluated based on multiclass predictions of 4D5 scFv variants binding to HER2 antigen. **(B, C, and D)** Meta learning and baseline machine learning model prediction performance is based on Matthew’s Correlation Coefficient (MCC) and plotted as a function of the number of training samples. FT refers to Fine-Tune Baseline. Points correspond to mean performance and shaded regions to 95 % confidence intervals across 3 random seeds. Performance curves plotted for meta sets consisting of 32 (black), 96 (purple), 288 (light blue), and 864 (gray) sequence-function pairs.

In contrast to the baselines, L2RW does not suffer performance loss under heavily downsampled training data with an MCC of approximately 0.8 with the CNN (Fig. 4E). Furthermore, performance rises to 0.9 as training and meta sizes are increased. MLC performs worse than the Fine-Tune baseline with small meta sets (Fig. 4D). This is expected as it is dependent on the source and extent of the noise as well as the nature of the algorithm: The meta label correction network requires a certain number of training examples and clean meta examples before it is able to begin correctly identifying and modifying noisy training data. Transformer MLC models surpassed the baseline models when trained with larger meta sets and above and with training sizes of 10% of full data and larger. Finally, performance differences are observed between the synthetic and experimental tasks, potentially resulting from differences in the nature of the noise (e.g. random versus systematic noise).

### Meta learning with positive and unlabeled data

The endpoint of classical directed evolution workflows is typically a library of protein variants enriched for the function of interest (referred to here as ‘positive’). For certain screening assays, it can be time consuming (e.g. FACS), difficult, or infeasible to collect the true negative sequences (e.g. growth-based selection) required to train supervised learning models. However, deep sequencing can be performed on the initial unscreened library and the final enriched library of positives. Supervised learning on such data has substantial challenges, as the unscreened library (referred to here as ‘unlabeled’) contains both positives and negatives. As an alternative, the method of Positive and Unlabeled Learning (PUL) seeks to learn from exactly this type of data. PUL has been successfully applied to many fields (Elkan and Noto, 2008; Bekker and Davis, 2020), including recently to protein sequence-function data to analyze the impact of amino acid substitutions on protein function and to design highly stable enzymes (Song *et al*., 2021). PUL methods are quite diverse, but often consist of either a two-step or a biased learning approach (Bekker and Davis, 2020). Although there are many varieties, the two-step approach generally seeks to: (i) estimate reliable negatives or the probability a positive example will be labeled and (ii) train a traditional classifier based on the estimation. Biased learning seeks to treat the unlabeled data as a noisy negative set and seeks to address the noise during learning. We hypothesize that meta learning may be beneficial in a PUL setting as it naturally falls under the latter category. For comparison, we introduce two additional baselines from the two-step PUL approach (described below).

#### 5A12 VEGF Synthetic PUL

As a proof-of-concept, we generated synthetic Positive Unlabeled (PU) training sets from deep sequencing data of the 5A12 antibody library following enrichment for High or Low/Non-binding to VEGF. The positive sets contain *n* sequences, consisting of enriched high binders. The unlabeled sets contain (*n* - α*n*) negatives and α*n* positives, with α varying from 0.1 to 0.8 (Fig. 5A). In this context, α can be thought of as the fraction of synthetic noise spiked into the negative set. The unlabeled set is treated as a noisy negative set during training of meta learning and traditional machine learning baseline algorithms. Two PUL baselines are added for comparison: PUDMS (Song *et al*., 2021) and ElkaNoto (Elkan and Noto, 2008). PUDMS is a PUL statistical framework recently adapted to protein sequence-function data from DMS. Two versions of PUDMS are tested, assuming either a linear (Order 1) or pairwise (Order 2) sequence-function mapping. ElkaNoto is a two-step approach that first trains a base classifier to estimate the probability a positive sample will be labeled via the label mechanism (FACS and deep sequencing in our case) and then trains a traditional classifier to predict positives and negatives. Two versions of the ElkaNoto approach are implemented as baselines, Random Forest and Logistic Regression. In general, the Fine-Tune and ElkaNoto baselines perform well, even with high α, when enough training data is available (Fig. 5 F,D,E). PUDMS is effective at maximizing the ratio of true positives to false positives, but scores inconsistently on the positive negative classification problem, with varying results for different data set sizes and performance decreasing with increasing α (Fig. 5 B,C). Traditional machine learning models and meta learning-aided Logistic Regression and Multilayer Perceptron models perform well on the task, however, the CNN and transformer models perform best in most cases for the synthetically generated PUL experiments (Fig. 5 G,H, Fig. S4, Fig. S5).

**Fig. 5.**
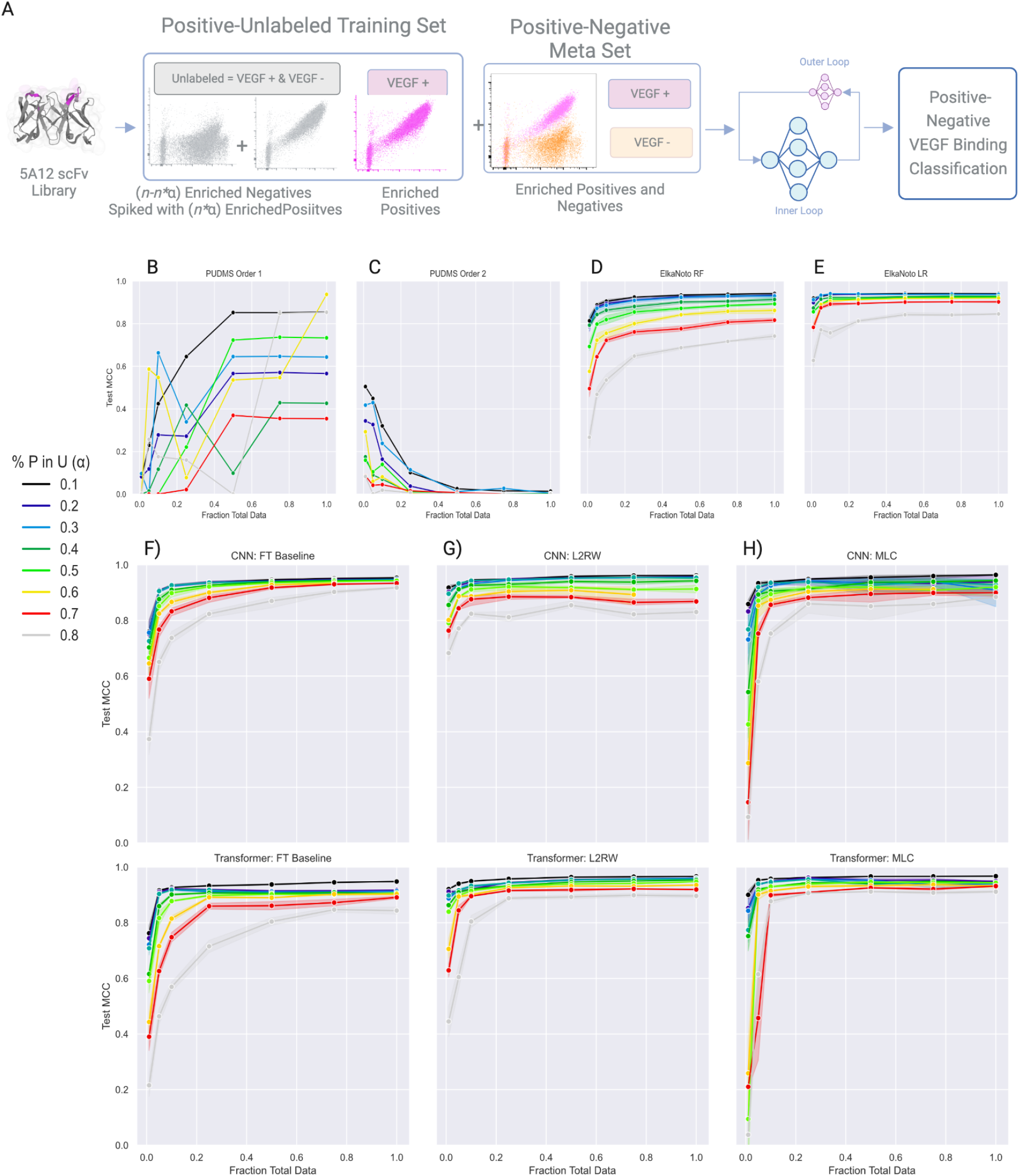
Meta learning applied to supervised machine learning models trained from yeast antibody (scFv) sequence data with synthetic Positive and Unlabeled data. **(A)** Schematic representation of machine learning task. The training, test, and meta data sets consist of deep sequencing of the 5A12 libraries following the final round of FACS (Fig. S2). Class labels refer to 5A12 variant populations with High- (positive), and Low/Non-binding (negative) to VEGF antigen. To create the unlabeled set, positives are synthetically mixed in with negatives at fraction α of the total unlabeled data set size. The meta and test set data remain unaltered, consisting of correctly labeled positive and negative sequence-function-pairs. Models are evaluated based on binary (positive/negative) binding classification predictions of 5A12 scFv variants to VEGF antigen. **(B-H)** Meta learning and baseline machine learning model prediction performance is based on Matthew’s Correlation Coefficient (MCC) and plotted as a function of the number of training samples. FT refers to Fine-Tune Baseline. Points correspond to mean performance and shaded regions to 95 % confidence intervals across 3 random seeds. A meta data set of 96 sequence-function pairs was used for all models.

#### Experimental PUL

The framework is next applied to experimentally relevant PU data: deep sequencing of the unscreened 5A12 antibody library is assigned as the unlabeled set and deep sequencing of the final round of enrichment for high binding to VEGF is assigned as the positive set. A range of meta set sizes are tested (32, 96, 288, 864 sequences). Each meta set is class balanced with an equal number of positives and negatives from their respective final enrichment rounds. Test sets are taken from the clean, final-round enrichments (Fig. 6A).

**Fig. 6.**
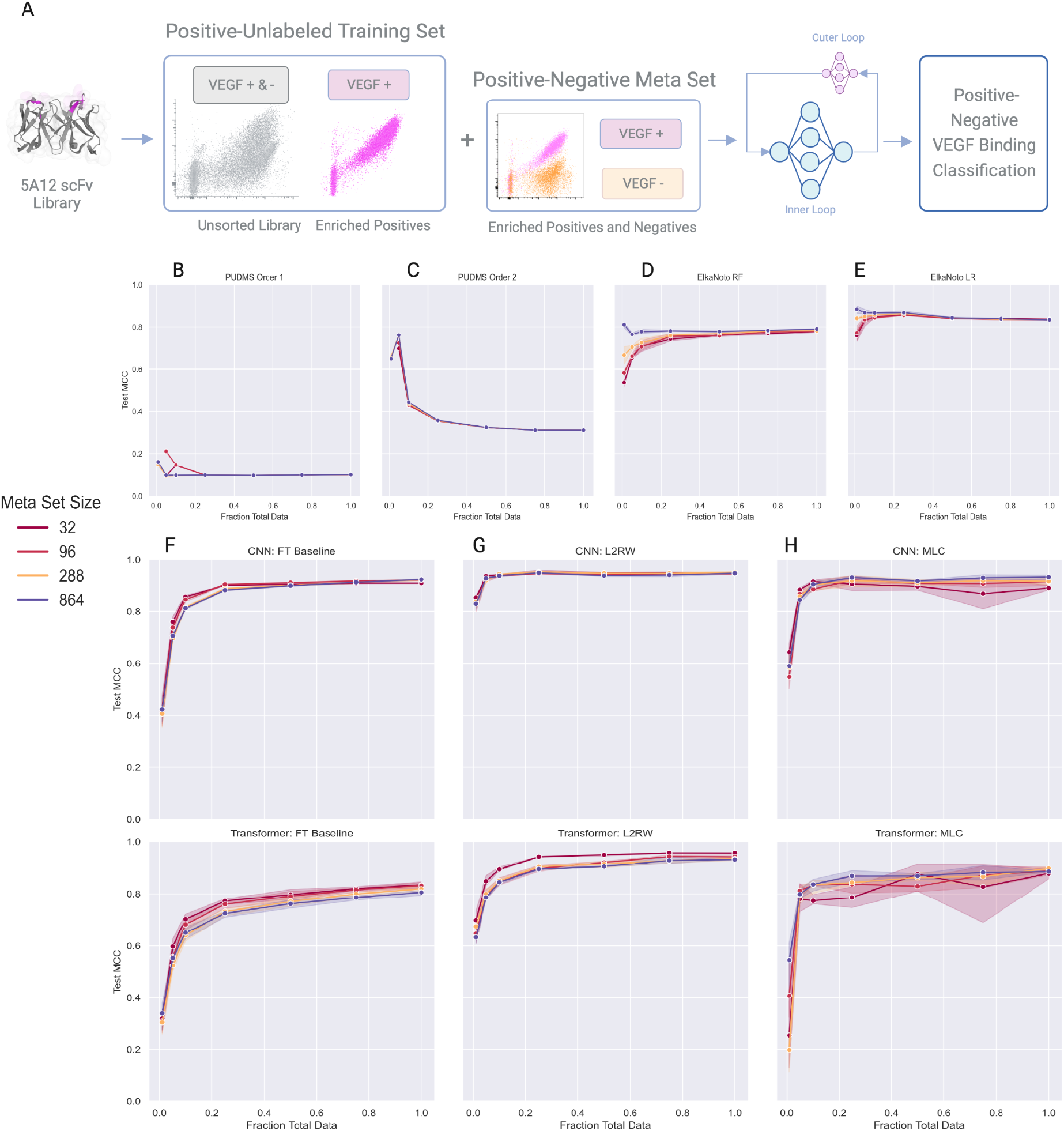
Meta learning applied to supervised machine learning models trained from yeast antibody (scFv) sequence data with experimental Positive and Unlabeled data. **(A)** Schematic representation of machine learning task. The training set consists of deep sequencing of i) positive (High-binding to VEGF antigen) 5A12 variants following the final round of FACS and ii) unlabeled 5A12 variants (both High- and Low/Non-binding to VEGF) from the unscreened library (Fig. S2). Test and meta data sets consist of deep sequencing of i) positive and ii) negative (Low/Non-binding to VEGF antigen) 5A12 variants following the final round of FACS. Models are evaluated based on binary (positive/negative) binding classification predictions of 5A12 scFv variants to VEGF antigen. **(B-H)** Meta learning and baseline machine learning model prediction performance is based on Matthew’s Correlation Coefficient (MCC) and plotted as a function of the number of training samples. FT refers to Fine-Tune Baseline. Points correspond to mean performance and shaded regions to 95 % confidence intervals across 3 random seeds. Performance curves plotted for meta sets consisting of 32 (dark red), 96 (light red), 288 (orange), and 864 (black) sequences.

The experimental PUL results generally mirror trends from synthetic PUL. PUDMS again demonstrates the ability to maximize the true positive to false positive ratio and scores inconsistently on the positive negative classification problem (Fig. 6 B,C). The ElkaNoto and Fine-Tune baselines perform well (Fig. 6 D,E,F). Traditional machine learning models and meta learning-aided Logistic Regression and Multilayer Perceptron models also perform well (Fig. S4, Fig. S5). However, the CNN and transformer models trained with L2RW and MLC typically outperform the baselines across meta set sizes. Meta learning trends are similar to the other tasks, e.g. MLC can be unstable with smaller meta sets and L2RW performs well with low training data (Fig. 6 F,G,H). Similar to the 4D5 tasks, differences between the synthetic and experimental PUL tasks could potentially stem from differences in the nature of the noise (e.g. random versus systematic noise).

### Learning multi-antigen classifiers from mostly single-antigen data

Binding to multiple antigens/targets is often a consideration in antibody drug discovery and development, e.g. to prevent off-target binding or to enable multi-target binding or cross-reactivity (Makowski *et al*., 2022; Saksena *et al*., 2022). Protein engineering campaigns of this type typically involve screening a library against multiple targets either in series or in parallel. Learning multi-target classifiers from largely single-target data could reduce overall screening burden, requiring screening only one target to completion and collecting labels for the remaining targets for a small number of variants. Towards this end we leveraged the multi-specificity of the 5A12 antibody, as it can bind with high affinity to both VEGF and Ang2 antigens. We constructed a representative task for the meta learning framework that seeks to predict Ang2 binding using a large training set of sequences labeled only for their ability to bind VEGF plus a small meta set with labels for both VEGF and Ang2 (i.e. present in both VEGF and Ang2 deep sequencing). We frame the task as four-class classification: (i) VEGF-positive and Ang2-positive (double positive), (ii) VEGF-positive and Ang2-negative, (iii) VEGF-negative and Ang2-positive, or (iv) VEGF-negative and Ang2-negative (double negative). Building on the experimentally determined VEGF binding label, we randomly assign half of sequences a positive label for Ang2 and the other half a negative Ang2 label, thereby introducing a high level of noise into the training set. Meta sets are batched from the set of experimentally confirmed sequences with labels for both VEGF and Ang2 antigens. Finally, as in previous tasks, we train with a noisy training set plus a clean meta set and test on a large, clean test set of variants experimentally screened for binding to both targets (present in deep sequencing following the final round of FACS for both VEGF and Ang2 screens).

Learning for this task appears more difficult than other experimental tasks, likely due in part to the large amount of noise introduced into the training sets during preprocessing (Fig. 7). The Fine-Tune baselines perform poorly for both CNN and transformer models (Fig. 7B). In contrast, L2RW and MLC yield reasonable performance at meta set sizes of 288 and 864 (Fig. 7 C,D). Although there is room for improvement, these results are encouraging as it demonstrates the potential of meta learning to reduce screening burden for this case using a relatively small number of experimentally validated sequences.

**Fig. 7.**
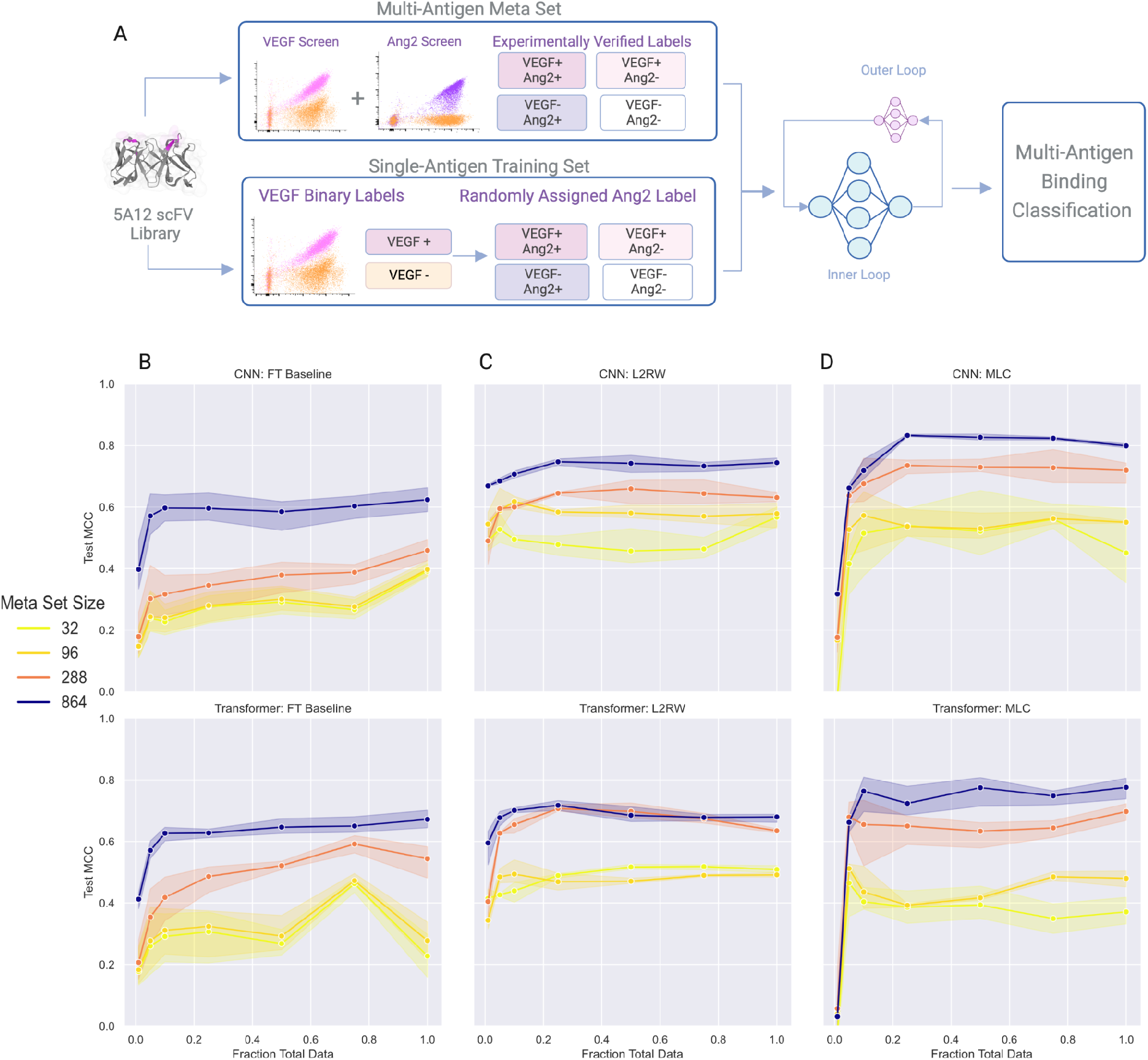
Meta learning applied to supervised machine learning models trained to predict multi-antigen binding classification using yeast antibody (scFv) sequence data with largely single-antigen binding classification labels. **(A)** Schematic representation of machine learning task. The training set is constructed from deep sequencing of 5A12 libraries following the final round of the VEGF FACS screen (Fig. S2). Positive (High-binding) and negative (Low/Non-binding) VEGF labels are retained and sequences are arbitrarily assigned an Ang2 binding classification. Test and meta sets consist of 5A12 variants with binding labels for both targets and are batched by combining deep sequencing from both VEGF and Ang2 final round FACS screens (Fig. S2, S3). **(B,C, and D)** Meta learning and baseline prediction performance (Matthew’s Correlation Coefficient) as a function of the number of training samples. FT refers to Fine-Tune Baseline. Points correspond to mean performance and shaded regions to 95 % confidence intervals across 3 random seeds. Performance curves plotted for meta sets consisting of 32 (bright yellow), 96 (dark yellow), 288 (red), and 864 (purple) sequences.

## Discussion

Rapid growth in machine learning-guided protein engineering is already resulting in impressive feats including generative protein design (Shin *et al*., 2021; Linder *et al*., 2020), engineering antibody affinity and specificity (Makowski *et al*., 2022), and structure-based models to engineer enzymes (Lu *et al*., 2022). The high-throughput assays required to collect large, labeled phenotype-genotype data sets, however, are time consuming and can yield noisy and under labeled data. Although much work has focused on maximizing the power of limited data and resources, for instance through informed training set design (Wittmann *et al*., 2021) and the use of pre-trained protein sequence embeddings (Hie *et al*., 2020; Biswas *et al*., 2021), most supervised learning approaches have utilized clean, fully labeled data sets (Makowski *et al*., 2022; Mason *et al*., 2021; Taft *et al*., 2022). We hypothesized that a meta learning framework could address these shortcomings and have the potential to reduce experimental screening burden.

Towards this end, we generated and screened two yeast display scFv libraries and created five experimentally relevant tasks to protein engineering workflows including synthetic and experimental label noise, positive and unlabeled learning, and multi-antigen learning from largely single-antigen data. The data for each task include a large, noisy training set, a small, clean meta set (ranging from 32 to 864 sequences), and a large, clean test set. We find that the meta learning algorithms L2RW (Ren *et al*., 2018) and MLC (Zheng *et al*., 2021) perform well using both a CNN and a transformer across all tasks, typically outperforming baselines. Additionally, these algorithms demonstrate strong performance even when only a small fraction of training data is available. It is worth noting that the Fine-Tune baseline performs well when large training and meta data sets are available. These results agree with studies reporting the ability of deep networks to maintain performance under noise given enough training data is available (Rolnick *et al*., 2018; Jiang *et al*., 2020).

Furthermore, performance differences in meta learning algorithms and baseline approaches are observed between synthetic and experimental noise and under labeled settings, likely due to the different source and nature of the noise between the setups. For example, synthetic label noise selects sequences for label corruption at random. Experimental noise, in contrast, may be dominated by systematic noise; for example, the noise in deep sequencing of High-binders to HER2 from the first 4D5 library screening round may consist of more Low-binders than Non-binders, as the former two populations are more similar to one another. These results suggest that understanding the source and extent of noisy or under labeled data can aid in the development process.

As demonstrated through this proof-of-concept study, meta learning is a powerful tool that has the potential to reduce experimental screening burden, improve robustness, and learn from the often imperfect data resulting from high-throughput screening experiments. For example, it may not be necessary to go through extensive enrichment rounds or to collect a large set of true negatives. Instead, one may choose to leverage meta learning to proceed more quickly to the next iteration of mutagenesis or characterization steps in a directed evolution or protein engineering workflow. Furthermore, the collection of a small, highly trusted meta data set is experimentally feasible for a variety of settings. However, the performance increases and robustness come at a computational cost, sometimes requiring a threefold increase in training time resulting from the multiple gradient descent steps required per batch. The computational cost, as well as the strong performance of the Fine-Tune baseline for multiple tasks, demonstrate the importance of fit-for-purpose methodology selection, highlighting that complex methods may not always be necessary or appropriate.

It is worth noting that in addition to protein engineering, meta learning has the potential to be useful for a wide variety of tasks in computational biology. Many experiments in biology result in noisy or under labeled data. Often, it is also feasible to obtain small, clean, and fully labeled data sets. Potential applications of meta learning may include leveraging single-cell sequencing to improve learning from noisy bulk sequencing; using known protein crystal structures as clean data to support computational prediction of structures (e.g., AlphaFold2, RosettaFold (Jumper *et al*., 2021; Baek *et al*., 2021)) that are especially difficult to model (e.g., the flexible CDR loops of antibodies and T cell receptors); using high-throughput screens like yeast display to measure thermostability or affinity in bulk via FACS and collecting small data sets with fine-grained instruments like differential scanning calorimetry and biolayer interferometry, respectively. Finally, future work may focus on algorithmic improvement, optimization of meta set composition and size, and the incorporation of pretrained embeddings and or structural information to potentially reduce data requirements and improve performance.

## Author contributions

Conceptualization, M.M and S.T.R.; Investigation, M.M; Software, M.M; Supervision, S.T.R.; Funding Acquisition, S.T.R; Writing – original draft, M.M, and S.T.R.

## Acknowledgements

We thank Dr. Roy Ehling, Dr. Joseph Taft, Jakub Kucharczyk, Beichen Gao, and Thomas Bikias for valuable scientific discussions. We are grateful to Dr. Joseph Taft for the HER2 expression plasmid. Computing resources and support from ETH Zurich (Euler cluster) are gratefully acknowledged. We also gratefully acknowledge the ETH Zurich D-BSSE Single Cell Facility and Genomics Facility, including Dr. Mariangela Di Tacchio, Renan Antonialli, Dr. Christian Beisel, Elodie Burcklen, Ina Nissen, and Mirjam Feldkamp.

## METHODS

### Key Resources Table

| Reagent or Resource | Source | Identifier |
| --- | --- | --- |
| Flow cytometry staining reagents |  |  |
| PE anti-DYKDDDDK Tag Antibody | Biolegend | 637310 |
| Biotinylated human VEGF165 | Acro | VE5-H82Q0-200ug |
| Biotinylated human angiopoietin-2 | Sino Biological | 10691-H08S-B |
| Deposited Data |  |  |
| Raw and processed sequencing data | This study |  |
| Oligonucleotides |  |  |
| Degenerate oPools for antibody library construction | IDT | https://github.com/LSSI-ETH/meta-learning-for-protein-engineering |
| Wild type gene fragments | Twist | https://github.com/LSSI-ETH/meta-learning-for-protein-engineering |
| Recombinant DNA |  |  |
| HER2 expression plasmid | (Li *et al.*, 2004) | RRID:Addgene_16257 |
| pYD1-4D5 (wt) | This study | https://github.com/LSSI-ETH/meta-learning-for-protein-engineering; derived from RRID:Addgene_73447 |
| pYD1-5A12 (wt) | This study | https://github.com/LSSI-ETH/meta-learning-for-protein-engineering; derived from RRID:Addgene_73447 |
| Cell Lines |  |  |
| EBY100 | ATCC | Cat# MYA-4941 |
| Software and Algorithms |  |  |
| L2RW | (Ren *et al.*, 2018) | https://github.com/uber-research/learning-to-reweight-examples |
| MLC | (Zheng *et al.*, 2021) | https://github.com/microsoft/MLC |
| ANT | (Engqvist and Nielsen, 2015) | https://github.com/jwillis0720/ANT |
| BBDUK | Joint Genome Institute | https://jgi.doe.gov/data-and-tools/software-tools/bbtools/bb-tools-user-guide/bbduk-guide/ |
| PUDMS | (Song *et al.*, 2021) | https://github.com/RomeroLab/pudms |
| ElkaNoto | (Elkan and Noto, 2008) | https://github.com/pulearn/pulearn |
| ggseqlogo | (Wagih, 2017) | https://github.com/omarwagih/ggseqlogo |
| Data preprocessing, model training, and analysis code | This study | https://github.com/LSSI-ETH/meta-learning-for-protein-engineering |

### Resource Availability

#### Lead contact

Further information and requests for reagents and resources should be directed to and will be fulfilled by the lead contact, Sai T. Reddy.

#### Materials availability

Antibody mutagenesis libraries generated in this study will be made available on request to the lead contact with a completed Materials Transfer Agreement.

#### Data and code availability

The main data supporting the results in this study are available within the paper and its supplemental information. Raw deep sequencing data will be deposited following publication. Code used for data curation, analysis and visualization are available at: https://github.com/LSSI-ETH/meta-learning-for-protein-engineering.

Additional data files and code that supports the findings of this study is available from the corresponding authors upon reasonable request.

#### Experimental Model and Subject Details

Saccharomyces cerevisiae EBY100 with yeast display (Boder and Wittrup, 1997) pYD1 plasmid were cultured 24-48 hours in a 250 rpm shaking incubator at 30 C in SD-UT medium (20 g/L glucose, 6.7 g/L yeast nitrogen base without amino acids, 5.4 g/L Na2HPO4, 8.6 g/L NaH2PO4·H2O and 5 g/l casamino acids). Induction was performed by incubating at 30 C (4D5) or 20 C (5A12) for 1 day in SG-UT induction medium (SD-UT with 20 g/l galactose substituted for glucose).

#### Method details

##### Yeast display scFv library design and transformation

Wild type 4D5 and 5A12 VH and VL scFv gene fragments were purchased from Twist Bioscience and transformed into the BamHI digested pYD1 plasmid via homologous recombination (Boder and Wittrup, 1997; Chao *et al*., 2006) using Frozen-EZ Yeast Transformation II (Zymo Research, T2001). 4D5 scFv was transformed in a HA-VL-218-VH-FLAG orientation and 5A12 scFv as HA-VH-218-VL-FLAG. HA and FLAG refer to the protein epitope tags and 218 to the linker.

Combinatorial mutagenesis scFv libraries were rationally designed from deep mutational scanning data (Mason *et al*., 2021; Koenig *et al*., 2015) and PDB crystal structures 1N8Z (Cho *et al*., 2003), 4ZFF, 4ZFG (Koenig *et al*., 2015) for 4D5-HER2, 5A12-VEGF, and 5A12-Ang2 respectively. Primers were designed and ordered from IDT with the desired amino acid degeneracy with the aid of the Python package ANT (Engqvist and Nielsen, 2015) to optimize the number of function-retaining mutations and are provided in Supplementary Note 1. Library inserts were synthesized via PCR using the wild-type scFv as template in a manner retaining at least 30 bp homology to the pYD1 backbone at both 5’ and 3’ ends. Libraries were cloned by homologous recombination using 2-4 μg template DNA and BamHI digested (linearized) pYD1 plasmid per 300μL EBY100 in a 2mm electroporation cuvette according to (Benatuil *et al*., 2010). The number of transformants were determined to be 3 x 10^7^ (4D5) and 9 x 10^7^ (5A12) via replicate SD-UT agar dilution plates.

##### Recombinant HER2 Expression and Fluorophore conjugation

HIS-tagged HER2 (Addgene #16257) was expressed using the Expi293™ expression system (Thermo Fisher, A14635) according to the manufacturer’s instructions. Briefly, 30 mL Expi293 cultures were transfected with HER2 expression plasmid and the supernatant harvested 5-7 days later via centrifugation at 300 G for 5 minutes followed by filtration (Steriflip 0.22mm Merck, SCGP00525). HER2 was then purified from supernatant as previously described (Vazquez-Lombardi *et al*., 2018). In short, supernatant was loaded to a TALON metal affinity (Clontech, 635502) gravity column, washed twice with PBS, and eluted with TALON elution buffer. Elution fractions were analyzed via SDS-PAGE and protein concentration quantified via Nanodrop 200c for A280 nm absorption. Protein-containing fractions were pooled and buffer exchanged with SnakeSkin™ dialysis tubing (10 MWCO, Pierce, 68100) followed by additional dialysis and concentration with Amicon Ultra-15 50 kDa centrifugal units (Merck, UFC905008). Purified protein was conjugated to AlexaFluor647 (Thermo Fisher, A3009) according to the manufacturer’s instructions.

##### Flow Cytometry Screening of Yeast Display Libraries

scFv libraries were cultured and induced as described above. Following induction, cells centrifuged at 8000 G for 2 minutes and washed with 1 mL wash buffer (Dulbecco’s PBS+ 0.5% BSA + 0.1% Tween20 + 2 mM EDTA). A one-step stain was used for the 4D5 library and a two-step stain used for 5A12. For 4D5, staining was performed for 30 minutes at 4 C with both 70 nM AlexaFluor647-conjugated HER2 and 1ng/μL anti-DYKDDDDK (anti-FLAG) Tag Antibody-PE (Biolegend, 637310). The 5A12 library was first incubated with either 0.5 nM VEGF (Acro, VE5-H82Q0-200ug) or 12 nM Ang2 (Sino Biological, 10691-H08S-B) plus Streptavidin-AlexaFluor647 (Bioloegend, 405237) for 30 minutes at 4 C. Following the fist stain, cells were centrifuged at 8000 G for 30 seconds and washed once before incubation with 1ng/μL anti-FLAG-PE for 30 minutes at 4 C. Following staining, cells were washed twice and kept on ice and away from light until sorting.

scFv expressing (FLAG+) cells in the 4D5 library were sorted by FACS (BD Aria Fusion) first into HER2 binding and non-binding fractions, then into three populations based on AlexaFluor647-conjugated HER2 Mean Fluorescent Intensity: High, Low, and Negative. scFv expressing (FLAG+) cells in the 5A12 library were sorted by FACS (BD Aria Fusion) into High binding (VEGF+ or Ang2+) or Low/Non-binding (VEGF- or Ang2-) populations. Sorted cells were cultured in SD-UT medium for 16-36 hours at 30 C. Induction and sorting was repeated for multiple rounds until the desired populations showed purity.

##### Deep Sequencing of scFv Libraries

Plasmid DNA encoding the scFv variants was isolated according to the manufacturer’s instructions (Zymo Research D2004). Variable regions of the scFv were amplified using custom primers. Illumina Nextera barcode sequences were then added in a second PCR amplification step, thereby facilitating high-throughput sequencing runs. Populations were pooled at the desired ratios and sequenced using the Illumina 2 x 250 PE protocol of the MiSeq Instrument.

##### Deep Sequencing Data Processing

Sequencing reads were quality trimmed and interleaved using BBDuk with a quality threshold 22. Mutagenesis regions were extracted, filtered for sequences with repeated reads adhering to the degenerate codons used for each library, and translated into amino acids using Python.

#### Quantification Statistical Analysis

All software developed for this study is found in the following link, at which a Python virtual environment is provided containing all packages and versions used: https://github.com/LSSI-ETH/meta-learning-for-protein-engineering.

##### Data Set Splitting and Truncation

Post deep sequencing processing with BBDUK and Python, all data preprocessing was carried out with Python (3.8.5) using pandas (1.5.0) and numpy (1.19.2). Following deep sequencing preprocessing as described above, amino acid sequences were assigned class labels (e.g. for 4D5 experiments, High-: 2, Low-: 1, Non-binding: 0). Sequences present in more than one population are removed. Levenshtein Distance, or Edit Distance (ED) from the consensus sequence of the data set was calculated for all sequences. An ED cutoff was used to separate the training and meta sets from the testing set in a manner so as to maximize the size and class balance in both the training and test sets (Table S5). An ED threshold of 6 (ED_6_) is applied to the 4D5 synthetic and experimental noise and 5A12 experimental PUL tasks. ED_5_ is applied to the 5A12 synthetic PUL and 5A12 Multi-Target tasks. Processed data set sizes are available in Table S5. Models are trained using training and meta sequences below the cutoff and tested on sequences above the cutoff. To mirror the greater sequence distribution, meta sets are batched from clean data below the cutoff using the sklearn (0.24.2) train_test_split function with ‘stratify’ option based on ED. Training and meta sets are concatenated for PUL and traditional machine learning baselines. Splitting training and testing via ED also resembles real-world workflows, in which models are trained with a limited number of mutations and used to extrapolate to a larger sequence space (Dallago *et al*., 2021).

To determine the impact data quantity has on model performance, training sets were truncated into separate subsets of 1%, 5%, 10%, 25%, 50%, 75%, and 100% of the full training data while keeping class balance ratios consistent and meta sets unmodified. To mirror the sequence distribution of the full data, truncation was performed using the sklearn (0.24.2) train_test_split function with ‘stratify’ option and truncated sets were stratified based on ED to the consensus sequence.

##### Data Visualization

Sequence logo plots (Fig. 1) were produced using the R package ggseqlogo (Wagih, 2017). Python (3.8.5) was used to compute machine learning metrics using scipy (1.9.2) and torchmetrics (0.7.2) and visualizations were generated with the matplotlib (3.4.3), seaborn (0.11.1), and numpy (1.19.2) packages.

##### Machine Learning Models And Computational Methods

###### Transformer Model

A modified transformer (Vaswani *et al*., 2017) was created with the following architecture. Protein sequences were categorically encoded as input to an embedding layer with dimension 32 followed by positional encoding injection and connected to a transformer encoder layer with 2 attention heads, each with hidden dimension 128. The encoder output was flattened to a linear layer with output dimension 512, followed finally by a linear layer with output dimension 1. Rectified Linear Unit (ReLU) was applied as the activation function and a dropout of 0.3 was used throughout the network.

###### CNN Model

One-hot encoded protein sequences were fed to the CNN as input. The network begins with two blocks, each consisting of 1D convolution, batch normalization, and max pooling layers. Both 1D convolution layers used a kernel width of 3 with 64 filters in the first convolution layer and 32 filters in the second. The output of the second block was flattened to a linear layer with 512 nodes and a dropout of 0.3, and final mapping to a linear layer with output dimension 1. ReLU was applied as the activation function throughout the network.

###### Model Training

Models were coded and trained in Python (3.8.5) using Pytorch (1.11.0) (Paszke *et al*., 2019) with CUDA (11.3). Cross Entropy was used as the loss function and the Matthews Correlation Coefficient (MCC), which ranges from −1 to 1, was selected as the performance metric due to its appropriateness for balanced, unbalanced, binary and multiclass tasks (Chicco and Jurman, 2020). Stochastic gradient descent (SGD) with momentum of 0.9 was used as the optimizer. A mini-batch size of 32 was used. Models were trained using the ETH Zurich Euler Cluster with 1 GPU (Nvidia GTX 1080, GTX 1080 Ti, or V100) and a requested 16 GB memory.

###### Deep Learning Methods

L2RW (Ren *et al*., 2018) was implemented using the Python package Higher (0.2.1) (Grefenstette *et al*., 2019). MLC was implemented as described in (Zheng *et al*., 2021) with a meta network embedding dimension 128 and hidden dimension of 64 and the k-step lookahead of 1. Training was carried out for 50 epochs. To mirror real-world experimental conditions and data availability, early stopping with patience was tested, but not implemented due to difficulties assessing performance on a validation set consisting of noisy or under labeled data. It should also be noted that in a noisy or under labeled learning setting, training loss and metrics are often unreliable indicators of true performance, further complicating matters. The Fine-Tune baseline trains a traditional classifier to completion then performs an additional 10 epochs of fine-tuning training on the meta set only. The Meta-Only baseline trains a traditional classifier on only the meta set.

###### Positive-Unlabeled Learning Baselines

PUDMS was implemented in R (4.1.3) according to (Song *et al*., 2021) for both Order 1 and Order 2 models. ElkaNoto (Elkan and Noto, 2008) was implemented via the Python pulearn library (https://pulearn.github.io/pulearn/). Random Forest and Logistic Regression ElkaNoto classifiers were implemented using sklearn (version 0.24.2). Random Forest was implemented with 200 trees and Logistic Regression with default parameters.

###### Meta Learning-Aided Logistic Regression and Multilayer Perceptron

Logistic Regression and Multilayer Perceptron (MLP) baselines were written in Pytorch (1.11.0) (Paszke *et al*., 2019) with CUDA (11.3) and trained in the same manner as the CNN and transformer. Both models take one-hot encoded protein sequences as input. Logistic Regression was implemented with default parameters. The MLP consists of a two-layer feed forward network: One-hot encoded sequences are flattened to a linear layer of dimension 256, followed by a second linear layer of dimension 512, before softmax output. Dropout of 0.3 and ReLU activation was applied throughout the network.

###### Traditional Machine Learning Model Baselines

Random Forest, Naive Bayes, and Linear Support Vector classifiers were implemented using sklearn (version 0.24.2). Random Forest was implemented with 200 trees and Linear Support Vector and Naive Bayes classifiers with default parameters.

