## Supplementary Material for "Meta Learning Improves Robustness and Performance in Machine Learning-Guided Protein Engineering"

### Supplementary Material for
### Meta Learning Improves Robustness and Performance in Machine Learning-Guided Protein Engineering

Authors: Mason Minot, Sai T. Reddy

Affiliation: ETH Zurich, Department of Biosystems Science and Engineering, Basel, 4058, Switzerland

**Supplementary Note 1. Degenerate Primers For scFv Library Generation**

4D5 CDRH3 degenerate reverse primers:

- AACTCATTTATCGTCGTCATCTTTATAATCGGATCCTGAGGAGACGGTGACCAGGGTTCCCTGGCCCCAGTAGTCCATCDSGTAGAAGCCRNNRNNMNNCCATCTGCTACAGTAATACAC
- AACTCATTTATCGTCGTCATCTTTATAATCGGATCCTGAGGAGACGGTGACCAGGGTTCCCTGGCCCCAGTAGTCCATCATGTAGAAGCCRNNRNNMNNCCATCTGCTACAGTAATACAC

4D5 CDRL3 degenerate reverse primers:

- CCTTGGTCCCTTGGCCGAAACCGGGAGGRRAMNNGWAGAACTGTTGACAGTAGTAAGTTG
- CCTTGGTCCCTTGGCCGAAGKTGGGAGGRRAMNNGWAGAACTGTTGACAGTAGTAAGTTG
- CCTTGGTCCCTTGGCCGAAACCGGGAGGCRTMNNGWAGAACTGTTGACAGTAGTAAGTTG
- CCTTGGTCCCTTGGCCGAAGKTGGGAGGCRTMNNGWAGAACTGTTGACAGTAGTAAGTTG
- CCTTGGTCCCTTGGCCGAAACCGGGAGGRRAMNNGWARTRCTGTTGACAGTAGTAAGTTG
- CCTTGGTCCCTTGGCCGAAGKTGGGAGGRRAMNNGWARTRCTGTTGACAGTAGTAAGTTG
- CCTTGGTCCCTTGGCCGAAACCGGGAGGCRTMNNGWARTRCTGTTGACAGTAGTAAGTTG
- CCTTGGTCCCTTGGCCGAAGKTGGGAGGCRTMNNGWARTRCTGTTGACAGTAGTAAGTTG

5A12 CDRH2 degenerate reverse primers:

- CCTTTCACGGAATCAGCGTAMNNMNNMMWGCCCCCGKYTRGAGTAATTCCAGCCACCCAT
- CCTTTCACGGAATCAGCGTAMNNMNNMMWGCCCCCCATTRGAGTAATTCCAGCCACCCAT

5A12 CDRL1 degenerate reverse primers:

- TAGCTTTGGGGCTTTGCCTGGCTTCTGTTGGTACCAGBTTACTCCMNNMYTGGATAAMNN
- TAGCTTTGGGGCTTTGCCTGGCTTCTGTTGGTACCAGBTTACTCCMNNCGGGATAAMNN
- TAGCTTTGGGGCTTTGCCTGGCTTCTGTTGGTACCATGCTACTCCMNNMYTGGATAAMNN
- TAGCTTTGGGGCTTTGCCTGGCTTCTGTTGGTACCATGCTACTCCMNNCGGGATAAMNN

**Table S1. 4D5 Sort Statistics**

| Population | Enrichment Round | Cells Collected |
|---|---|---|
| HER2 Positive / FLAG+ | 0 | $1.9 \times 10^7$ |
| HER2 High / FLAG+ | 1 | $1.3 \times 10^7$ |
| HER2 High / FLAG+ | 2 | $1.0 \times 10^7$ |
| HER2 High / FLAG+ | 3 | $9 \times 10^6$ |
| HER2 Low / FLAG+ | 1 | $9 \times 10^6$ |

| | | |
|---|---|---|
| HER2 Low / FLAG+ | 2 | $1.3 \times 10^7$ |
| HER2 Low / FLAG+ | 3 | $1.4 \times 10^6$ |
| HER2 Negative / FLAG+ | 0 | $1.8 \times 10^7$ |
| HER2 Negative / FLAG+ | 1 | $4.0 \times 10^7$ |
| HER2 Negative / FLAG+ | 2 | $6.0 \times 10^7$ |
| HER2 Negative / FLAG+ | 3 | $7.0 \times 10^7$ |

**Table S2. 5A12 VEGF Sort Statistics**

| Population | Enrichment Round | Cells Collected |
|---|---|---|
| VEGF+ / FLAG+ | 0 | $3.7 \times 10^7$ |
| VEGF+ / FLAG+ | 1 | $4.2 \times 10^7$ |
| VEGF+ / FLAG+ | 2 | $6.3 \times 10^7$ |
| VEGF+ / FLAG+ | 3 | $2.4 \times 10^7$ |
| VEGF- / FLAG+ | 0 | $3.2 \times 10^7$ |
| VEGF- / FLAG+ | 1 | $1.5 \times 10^7$ |
| VEGF- / FLAG+ | 2 | $6.9 \times 10^7$ |

**Table S3. 5A12 Ang2 Sort Statistics**

| Population | Enrichment Round | Cells Collected |
|---|---|---|
| Ang2+ / FLAG+ | 0 | $2.0 \times 10^7$ |
| Ang2+ / FLAG+ | 1 | $3.0 \times 10^7$ |
| Ang2- / FLAG+ | 0 | $2.0 \times 10^7$ |
| Ang2- / FLAG+ | 1 | $3.0 \times 10^7$ |

**Table S4. Deep Sequencing Statistics**

| Library | Screening Round | Population | Number Reads | Unique Amino |
|---|---|---|---|---|

| | | | | Acid Sequences |
|---|---|---|---|---|
| 4D5 | Initial Sort | Negative | $8.5 \times 10^6$ | $4.7 \times 10^5$ |
| | | Low | $9.0 \times 10^6$ | $4.7 \times 10^5$ |
| | | High | $4.5 \times 10^6$ | $1.2 \times 10^5$ |
| | First Sort | Negative | $6.3 \times 10^6$ | $6.3 \times 10^5$ |
| | | Low | $9.5 \times 10^6$ | $5.6 \times 10^5$ |
| | | High | $6.3 \times 10^6$ | $5.4 \times 10^5$ |
| 5A12 (VEGF) | Final Sort | Low/Negative | $4.9 \times 10^6$ | $2.6 \times 10^5$ |
| | | High | $7.5 \times 10^6$ | $3.6 \times 10^5$ |
| | Unscreened | Unlabeled | $5.5 \times 10^6$ | $3.7 \times 10^5$ |
| 5A12 (Ang2) | Final Sort | Low/Negative | $1.1 \times 10^7$ | $7 \times 10^5$ |
| | | High | $7.7 \times 10^6$ | $1.6 \times 10^4$ |

**Table S5. Processed Data Set Sizes Per Task**

| Task | Data Set | Number of Protein Sequences |
|---|---|---|
| 4D5 Synthetic Noise | Train | 105,813 |
| | Test | 117,657 |
| 4D5 Experimental Noise | Train | 242,721 |
| | Test | 48,078 |
| 5A12 VEGF Synthetic PUL | Train | 171,338 |
| | Test | 257,988 |
| 5A12 VEGF Experimental PUL | Train | 202,416 |
| | Test | 87,872 |
| 5A12 Multi-Target | Train | 135,272 |
| | Test | 25,439 |

*Note:* Meta sets are created for each data set in sizes of 32, 96, 288, and 864 sequences from clean / fully labeled data under the Edit Distance train/test split cutoff.

**Table S6. MetaSet-Only baseline results**

| Task | Base Model | Meta Set Size | MCC |
|---|---|---|---|
| 4D5 Synthetic Noise | CNN | 96 | 0.57 ± 0.01 |
| | Transformer | 96 | 0.37 ± 0.01 |
| | Logistic Regression | 96 | 0.23 ± 0.08 |
| | Multilayer Perceptron | 96 | 0.13 ± 0.13 |
| 4D5 Experimental Noise | CNN | 32 | 0.44 |
| | | 96 | 0.45 |
| | | 288 | 0.53 |
| | | 864 | 0.61 |
| | Transformer | 32 | 0.30 |
| | | 96 | 0.41 |
| | | 288 | 0.45 |
| | | 864 | 0.61 |
| | Logistic Regression | 32 | 0.20 ± 0.09 |
| | | 96 | 0.21 ± 0.09 |
| | | 288 | 0.24 ± 0.09 |
| | | 864 | 0.34 ± 0.07 |
| | Multilayer Perceptron | 32 | 0.13 ± 0.18 |
| | | 96 | 0.15 ± 0.18 |
| | | 288 | 0.13 ± 0.14 |
| | | 864 | 0.20 ± 0.23 |
| 5A12 VEGF Synthetic PUL | CNN | 96 | 0.54 |
| | Transformer | 96 | 0.43 |
| | Logistic Regression | 96 | 0.19 ± 0.07 |
| | Multilayer Perceptron | 96 | 0.27 ± 0.10 |
| 5A12 VEGF Experimental PUL | CNN | 32 | 0.38 |
| | | 96 | 0.51 |

| | | | |
|---|---|---|---|
| | | 288 | 0.54 |
| | | 864 | 0.61 |
| | Transformer | 32 | 0.36 |
| | | 96 | 0.43 |
| | | 288 | 0.49 |
| | | 864 | 0.69 |
| | Logistic Regression | 32 | 0.28 |
| | | 96 | 0.22 ± 0.17 |
| | | 288 | 0.26 ± 0.16 |
| | | 864 | 0.35 ± 0.14 |
| | Multilayer Perceptron | 32 | 0.23 ± 0.14 |
| | | 96 | 0.27 ± 0.14 |
| | | 288 | 0.31 ± 0.16 |
| | | 864 | 0.39 ± 0.13 |
| 5A12 Multi-Target | CNN | 32 | 0.32 |
| | | 96 | 0.32 |
| | | 288 | 0.41 |
| | | 864 | 0.53 |
| | Transformer | 32 | 0.23 |
| | | 96 | 0.22 |
| | | 288 | 0.30 |
| | | 864 | 0.47 |
| | Logistic Regression | 32 | 0.14 ± 0.11 |
| | | 96 | 0.13 ± 0.11 |
| | | 288 | 0.15 ± 0.12 |
| | | 864 | 0.35 ± 0.13 |
| | Multilayer Perceptron | 32 | 0.06 ± 0.02 |

| | | | |
|---|---|---|---|
| | | 96 | 0.08 ± 0.02 |
| | | 288 | 0.09 ± 0.05 |
| | | 864 | 0.26 ± 0.10 |

*Note:* Mean and standard deviation MCC reported across 3 seeds. Standard deviation less than ± 0.00 unless noted otherwise.

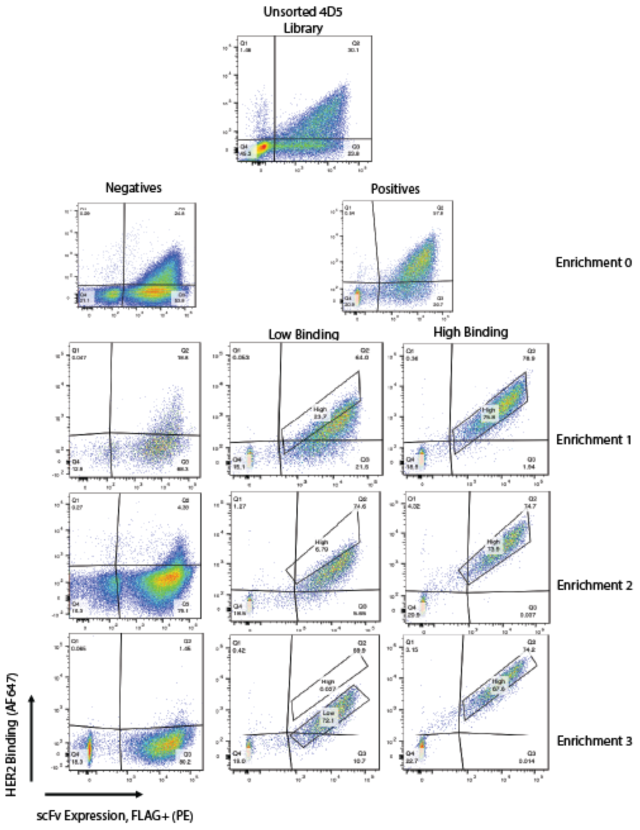


**Fig. S1** 4D5 Sort Summary. 4D5 antibody library FACS plots from initial library through final enrichment round. Y-axis corresponds to HER2 binding (AF647) and X-axis to scFv expression (FLAG+ / PE).

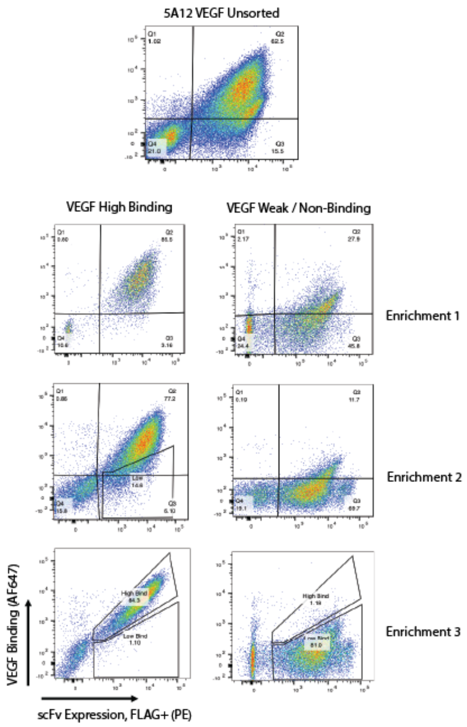


**Fig. S2** 5A12 VEGF Sort Summary. 5A12 antibody library FACS plots from initial library through final enrichment round. Y-axis corresponds to VEGF binding (AF647) and X-axis to scFv expression (FLAG+ / PE).

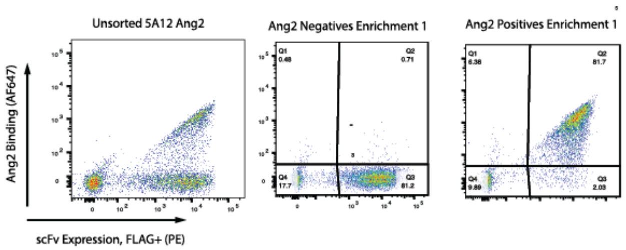


**Fig. S3** 5A12 Ang2 Sort Summary. 5A12 antibody library FACS plots from initial library through final enrichment round. Y-axis corresponds to Ang2 binding (AF647) and X-axis to scFv expression (FLAG+ / PE).

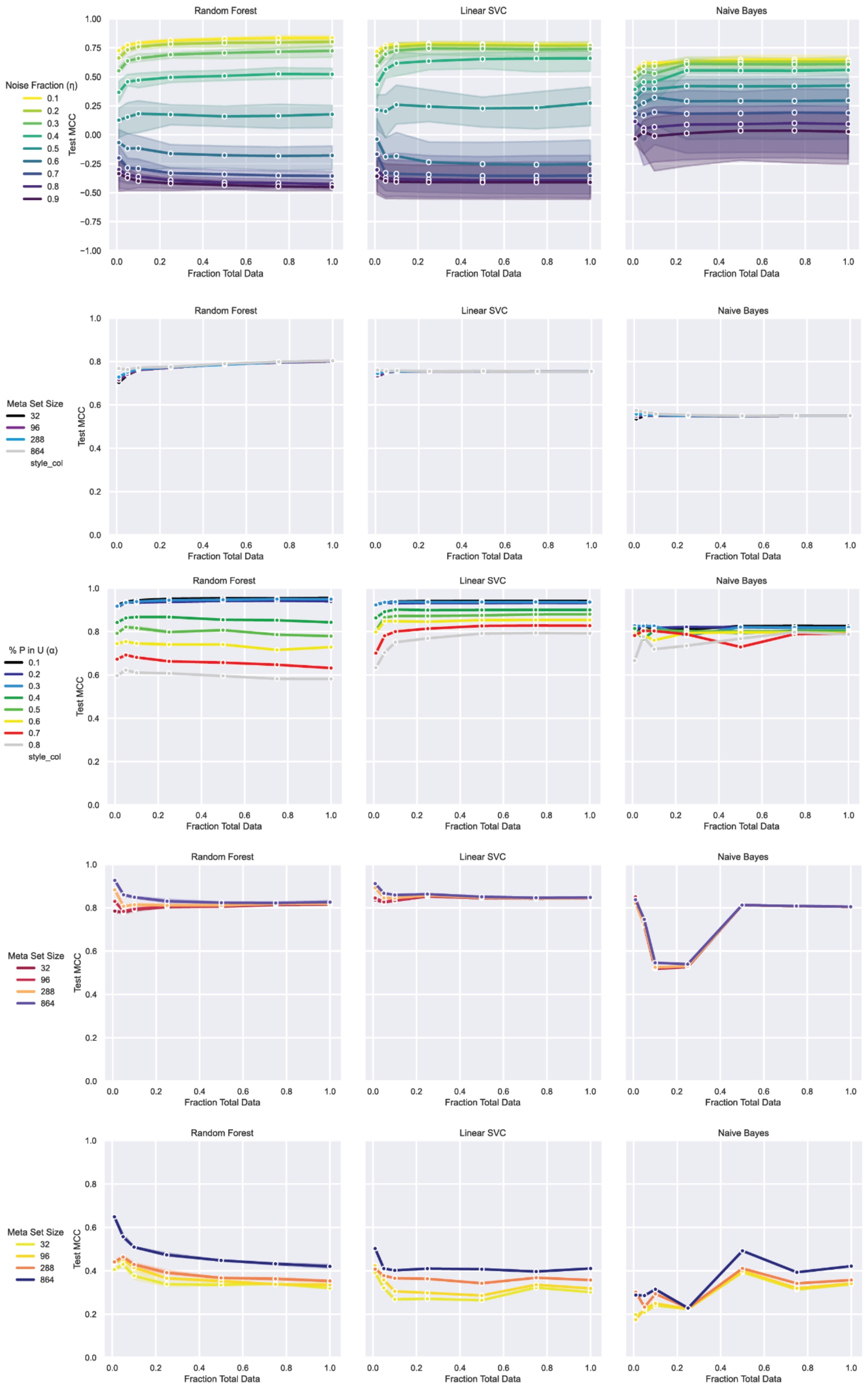


**Fig. S4** Traditional machine learning models evaluated on each task: Random Forest, Linear Support Vector Classifier (Linear SVC), and Naive Bayes. Prediction performance (Matthew's Correlation Coefficient, or MCC) as a function of the number of training samples. Points correspond to mean performance and shaded regions to 95 % confidence intervals across 3 random seeds.

**(A)** 4D5 synthetic noise task.
**(B)** 4D5 experimental noise task.
**(C)** 5A12 synthetic PUL task.
**(D)** 5A12 experimental PUL task.
**(E)** 5A12 multi-antigen task.

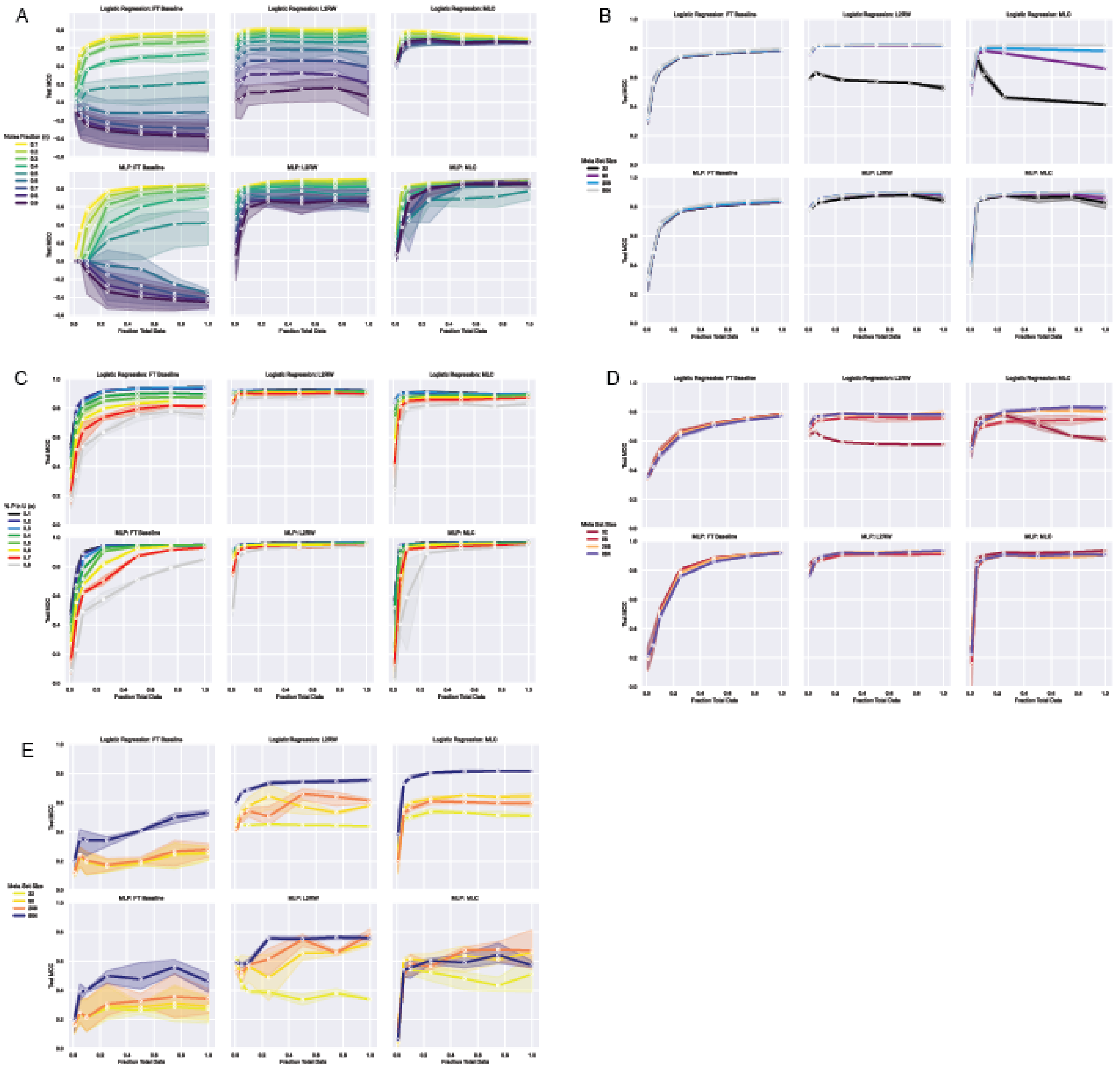

**Fig. S5** Meta learning algorithms evaluated on each task with Logistic Regression and Multilayer Perceptron Classifiers. Prediction performance (Matthew's Correlation Coefficient) as a function of the number of training samples. Points correspond to mean performance and shaded regions to 95 % confidence intervals across 3 random seeds.
**(A)** 4D5 synthetic noise task.
**(B)** 4D5 experimental noise task.
**(C)** 5A12 synthetic PUL task.
**(D)** 5A12 experimental PUL task.
**(E)** 5A12 multi-antigen task.
